# Neutralization Of SARS-CoV-2 Variants By A Human Polyclonal Antibody Therapeutic (COVID-HIG, NP-028) With High Neutralizing Titers To SARS-CoV-2

**DOI:** 10.1101/2022.01.27.478053

**Authors:** Derek Toth

## Abstract

Since the start of the COVID-19 outbreak the World Health Organization (WHO) has classified multiple SARS-CoV-2 Variants-of-Concern and Variants-of-Interest (VOC/VOI) with mutations in their Spike protein that increase transmissibility and/or reduce the effectiveness of vaccines and monoclonal antibody therapeutics. The emergence of these variants represents a significant health risk and highlights the need for additional COVID-19 therapeutics that maintain the ability to neutralize current, as well as future variants.

COVID-HIG (NP-028) is a polyclonal Anti-SARS-CoV-2 human Immunoglobulin purified from source human plasma screened for high antibody titers to SARS-CoV-2 antigens. COVID-HIG was previously evaluated in INSIGHT 013 clinical trial [NCT04546581] which was an international, multi-center, adaptive, randomized, double-blind, placebo-controlled trial of the safety, tolerability and efficacy of a single dose infusion (up to 400 mL) of Anti-Coronavirus Hyperimmune Intravenous Immunoglobulin (hIVIG) for the treatment of adult recently hospitalized COVID-19 patients (N=593). COVID-HIG is currently being evaluated for clinical efficacy in a Phase 3 placebo-controlled study INSIGHT 012 (NCT04910269) to compare the safety and efficacy of a single infusion of anti-COVID-19 hyperimmune immunoglobulin (hIVIG) versus placebo among adults with recently diagnosed SARS-CoV-2 infection who do not require hospitalization.

In the present study, in-vitro pseudovirus and live virus neutralization assays were used to assess the impact of SARS-CoV-2 variant spike mutations on neutralizing potency of COVID-HIG. These assays are valuable tools for monitoring the potential impact of variant mutations on efficacy of antibody therapeutics as well as vaccines/natural immunity.

To date, COVID-HIG (NP-028) has been shown to retain neutralizing potency against 20 full spike protein sequence SARS-CoV-2 pseudovirus variants including all currently classified VOC/VOI (Alpha, Beta, Gamma, Delta/Delta+, Eta, Iota, Kappa, Lambda, Mu as of Sept 2021) as well as 4 live virus variants (Alpha, Beta, Gamma, and Iota).

## Introduction

Since the start of the COVID-19 outbreak in late 2019, numerous variants of SARS-CoV-2 have emerged around the world (Zhou et al., 2020). The WHO in collaboration with partners, expert networks, national authorities, institutions, and researchers have been monitoring the evolution of SARS-CoV-2 since January 2020(WHO, 2021).

SARS-CoV-2 variants with characteristics that pose an increased risk to global public health have been classified by WHO as Variants of Interest (VOIs) and Variants of Concern (VOCs) so that global monitoring and research can be prioritized in response to the COVID-19 pandemic. In May 2021 the WHO COVID-19 reference library network established a nomenclature for each VOC/VOI using the Greek alphabet (e.g. Alpha, Beta, Gamma, Delta, etc.) to assist in identifying each and for communicating to the public.

The Alpha (B.1.1.7) variant was first to be classified as a VOC and was detected in the United Kingdom in October 2020. Since then, it has been reported in 174 countries around the world including 50 US states(Mullen et al., 2020). The Alpha variant is ~50% more transmissible than original strain and has the potential for increased severity based on hospitalizations and case fatalities (Horby et al., 2021) but with no reported significant impact on neutralization by EUA monoclonal antibody treatments, convalescent plasma, or post vaccination sera (Wang et al., 2021b).

The Beta (B.1.351) variant was first identified in South Africa on Dec 18, 2020 and has since been reported in 115 countries including 50 US states (Mullen et al., 2020). The Beta variant is ~50% more transmissible than original strain (Pearson et al., 2021) and is significantly less susceptible to neutralization by the monoclonal antibody treatment combination of bamlanivimab and etesevimab, as well as convalescent plasma and post-vaccination sera (Wang et al., 2021b). It has been reported in South Africa that the prevalence of the Beta variant is higher in young people with no underlying health conditions, and by comparison to other variants, more frequently results in serious illness in those cases (Mkhize, 2020).

The Gamma (P.1) variant was first detected on January 6, 2021 in four travelers returning to Tokyo after travel to Brazil. The Gamma variant was confirmed in Brazil on Jan 12, 2021 and has been detected in at least 83 countries around the world including 50 US states(Mullen et al., 2020). The Gamma variant has been reported to exhibit increased transmissibility in both young and old, higher viral load in infected individuals, significantly reduced susceptibility to neutralization by the monoclonal antibody treatment of bamlanivimab and etesevimab, as well as convalescent plasma and post-vaccination sera (Wang et al., 2021a). The Gamma variant has also been associated with more serious illness and higher mortality (Faria et al., 2021).

The Delta (B.1.617.2) variant was first detected in India in October 2020 and has since been reported in 167 countries around the world including 50 US states (Mullen et al., 2020). A Delta variant with an additional mutation (K417N) has also been identified and given the Delta+ designation. The K417N mutation found in the Delta+ variant is shared with the Beta and Gamma variants and may increase transmissibility, reduce the effectiveness of certain monoclonal antibody therapeutics as well as convalescent plasma and post-vaccination sera. The Delta/Delta+ variants have been responsible for a significant increase in global infections, including vaccine breakthrough infections (Acharya and Jamkhandikar, 2021), and are the predominant variants in most countries including the US as of Sept 21, 2021 (Fig 1).

**Figure 1:**
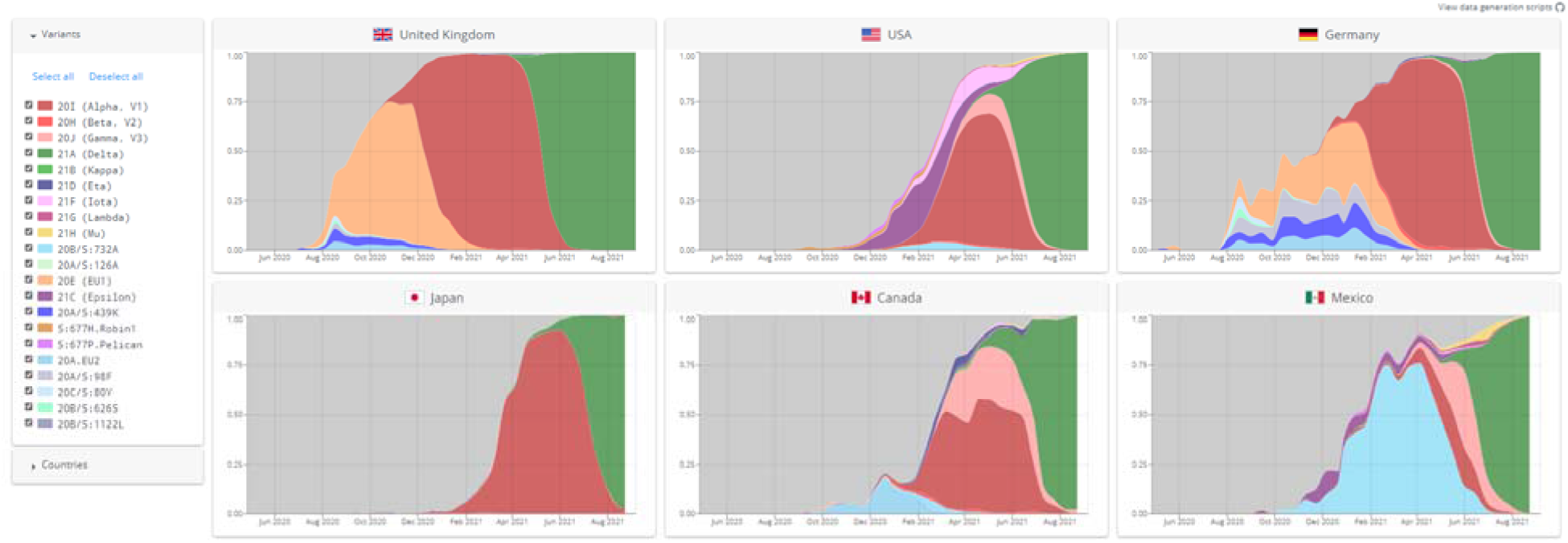
Graphical representation of proportion of sequenced SARS-CoV-2 variant groups in selected countries over time (as of Sept 21, 2021) The above figure is an unmodified screenshot from https://covariants.org/ and used under license https://creativecommons.org/licenses/by/4.0/

Even though natural immunity, vaccine induced immunity and monoclonal antibody therapeutics can be effective in preventing severe SARS-CoV-2 infection/re-infection, breakthrough infections from natural and vaccine immunity have been reported against VOC/VOIs (Gazit et al., 2021). In addition, antibody therapeutics (individual monoclonals/monoclonal cocktails and convalescent plasma) have been shown to have significantly reduced neutralizing efficacy to many of the SARS-CoV-2 VOC/VOI due mutations in the spike protein (Chen et al., 2021a; Chen et al., 2021b; Planas et al., 2021).

With the current circulating VOC/VOIs and the potential for more variants to arise in the future, it is important to evaluate SARS-CoV-2 antibody therapeutics for in-vitro neutralizing activity across multiple VOC/VOIs to help inform efficacy evaluations and protect populations at risk of COVID-19.

Emergent’s COVID-HIG, NP-028 was evaluated in a comparative profiling study initiated in October 2020 by the U.S. Government COVID-19 Response Therapeutics Research team. COVID-HIG along with 16 individual monoclonals, 6 monoclonal cocktails and 3 polyclonal clinical stage SARS-CoV-2 therapeutic antibodies were tested for neutralizing activity against 60+ pseudoviruses (Lusvarghi et al., 2021) and 4 live virus variants with single or multiple substitutions, including full set substitutions in the spike protein for current VOC/VOI’s (Alpha, Beta, Gamma, Delta/Delta+, Eta, Iota, Kappa, Lambda and Mu). In the published study, Emergent’s COVID-HIG, NP-028 is reported as pABs blinded code III.

### Natural Immunity, Vaccine Immunity, Passive Transfer Immunity of Antibody Therapeutics (convalescent plasma, monoclonals, polyclonal hyperimmune)

**Natural immunity** is developed when viral pathogens circulate and cause infections in populations. Individuals that recover, develop an adaptive cellular and polyclonal antibody humoral immune response to multiple epitopes of the pathogen. This adaptive cellular and humoral response helps clear the initial infection and prevents re-infections (or reduced severity), if reexposed.

**Vaccines** confer similar protection to natural immunity and are indicated for active immunization to prevent coronavirus disease 2019 (COVID-19) caused by severe acute respiratory syndrome coronavirus 2 (SARS-CoV-2). Most vaccines require several doses over multiple weeks before an individual develops an immune response that can prevent or reduce severity of infection. Additionally, vaccines may not be effective in high-risk populations with compromised immune systems due to advanced age, immune deficiency diseases, immunosuppression therapy and hematologic malignancies, etc. (Lee et al., 2021). Vaccines may also lose efficacy over time due to waning immunity, require boosters as new variants arise resulting in breakthrough infections, or new vaccines developed if previous vaccines are no longer effective against new variants (Gazit et al., 2021).

**Convalescent plasma (CP)** can be collected from individuals fully recovered from infection and used for passive immunotherapy. The efficacy of CP can vary significantly between donors as each individual has a unique humoral immune response. Each unit of CP will have varying titers of neutralizing antibody, different IgG isotype profile, variable antibody binding affinity and recognition of different virus proteins and epitopes. Antibody titers in convalescent plasma may also significantly decrease over time which may result in reduced or loss of efficacy (Lau et al., 2021). Depending upon variant exposure and individual immune response, convalescent plasma from some donors may cross-neutralize virus variants, whereas some may not (Schmidt et al., 2021). Therefore, convalescent plasma (CP) from recovered individuals should be collected early after symptoms resolve and tested for antibody titers to circulating variants prior to use to ensure potency.

**Monoclonals** can be developed for passive immunotherapy to prevent or treat viral infections using the traditional hybridoma approach or engineered from B-cells collected from convalescent patients. Monoclonals are well characterized, recognize single epitopes on viral antigens, have high potency and can be produced consistently over time. Two or more monoclonals that bind different epitopes can be combined to improve efficacy and reduce the potential for resistance from mutations found in virus variants. As monoclonals/monoclonal cocktails recognize 1-2 neutralizing epitopes on virus target proteins, only a few mutations in the target virus protein may result in complete loss of monoclonal efficacy. Also, as monoclonal/monoclonal cocktails are given prophylactically or for treatment, the selective pressure over time may drive viral mutations resulting in escape mutants and significantly reduced or complete loss of neutralizing efficacy (Harvey et al., 2021).

**Passive transfer immunotherapy** can provide immediate protection where antibodies to the virus contained in convalescent plasma, purified immunoglobulins (hyperimmunes) and monoclonal/monoclonal cocktails are administered for treatment and/or prevention of infection. Passive immunotherapy with polyclonal hyperimmunes has a had a long and successful history for treatment or prevention of many types of viral/bacterial infections. Currently there are seven human polyclonal immunoglobulin products that have been approved against various viral/bacterial infections, including respiratory infections like RSV (Tharmalingam et al., 2021). Once convalescent/vaccinated donor plasma is available, it can be screened for high titers to the target virus, pooled, purified and concentrated into IgG using established GMP manufacturing platforms. The plasma pool titers for each manufacturing run are carefully controlled to ensure suitable potency and tested using validated quality control assays prior to lot release. Since hyperimmunes are manufactured by combining plasma from multiple donors, each lot contains the combined polyclonal antibody profile of every individual plasma donor included in the manufacturing pool. This results in a broad range of antibodies that bind to many viral proteins/epitopes and is significantly broader than monoclonals/monoclonal cocktails as well as individual units of convalescent plasma. The broad range polyclonality of hyperimmunes should reduce the risk of losing potency to current and future variants and as new variants arise, hyperimmunes can adapt over time as plasma from donors who recover from variant infections can be screened for suitable titer and used to manufacture future lots. This concept is currently leveraged in the use of Intravenous Immune Globulin (IVIG) products for the prevention of infections in primary immune deficiency (PID) patients, where the diverse plasma donor population (>1000 donors) for IVIG products provides a high likelihood that it will contain antibodies to pathogens circulating in the population.

Currently there are several passive immunotherapeutic products in clinical trials to treat or prevent SARS-CoV-2 infection with some granted Emergency Use Authorization (EUA) or full FDA approval. Most of these passive immunotherapeutics are monoclonals or monoclonal cocktails that target the Spike protein (Receptor Binding Domain -RBD and/or N-Terminal Domain – NTD) and are based on the amino acid sequence from the original Wuhan isolate or derived from patients who recovered from infection early in the COVID-19 outbreak. Several monoclonals that are EUA-authorized and under development have demonstrated reduced efficacy against certain emerging SARS-CoV-2 variants (Chen et al., 2021a; Chen et al., 2021b; Planas et al., 2021). If these variants are to become predominant or continue to evolve to escape detection, monoclonal antibody therapeutics could become obsolete.

New vaccines and monoclonal antibodies can be developed to provide protection against these new variants, but new clinical trials and regulatory approval would likely be required before they would be widely available. Polyclonal antibodies in hyperimmunes are typically more robust and resistant to mutations and variants as they bind a diverse array of epitopes. Therefore, developing a polyclonal antibody option is critical to ensure effective antibody treatments are available long-term.

Emergent’s fully human polyclonal IgG antibody hyperimmune (COVID-HIG, NP-028) is manufactured from a pool of plasma from multiple donors screened to have high antibody titers to the Spike protein of SARS-CoV-2 and tested for neutralizing potency by validated in-vitro potency assays. COVID-HIG was previously evaluated in INSIGHT 013 clinical trial [NCT04546581] which was an international, multi-center, adaptive, randomized, double-blind, placebo-controlled trial of the safety, tolerability and efficacy of a single dose infusion (up to 400 mL) of Anti-Coronavirus Hyperimmune Intravenous Immunoglobulin (hIVIG) for the treatment of adult recently hospitalized COVID-19 patients (N=593). COVID-HIG is currently being evaluated in a phase 3 clinical trial OTAC INSIGHT 012 (NCT04910269): An International Multicenter, Adaptive, Randomized Double-Blind Placebo-Controlled Trial of the Safety, Tolerability, and Efficacy of Anti-Coronavirus Hyperimmune Intravenous Immunoglobulin (hIVIG) for the Treatment of Adults Outpatients in Early Stages of COVID-19 (Clinicaltrials.gov NCT04910269).

In this report un-blinded results for Emergent’s human polyclonal hyperimmune COVID-HIG (NP-028) are provided showing its ability to neutralize all 20 full sequence pseudovirus and 6 live virus variants tested in-vitro to date (table 1), including the Delta/Delta+ variants, which are currently the predominant variants driving the 4^th^ wave in the US and the world (Hodcroft, 2021). Additional SARS-CoV-2 variant data on Emergent’s COVID-HIG in comparison to other monoclonal/polyclonal antibody therapeutics can be found in (Lusvarghi et al., 2021) under blinded code III. Of note, the COVID-HIG lot used in this study was an early small-scale pilot with a plasma pool of only 19 donors. All the plasma used in this pilot lot was collected prior to May of 2020, which was relatively early in the outbreak and before VOC/VOI’s were circulating at significant levels in the donor population.

**Table 1:**
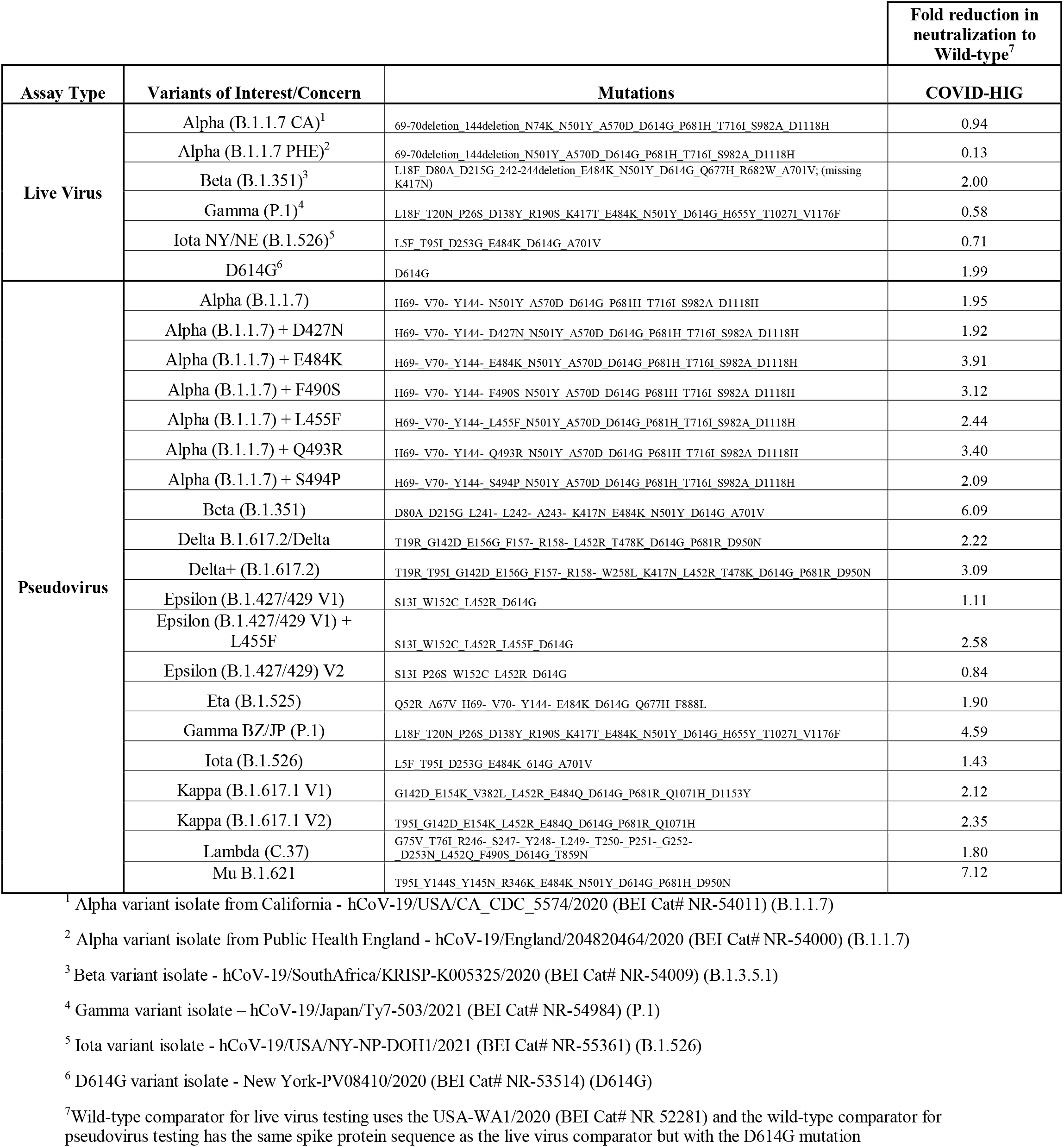

## Results

FDA in collaboration with NIAID published (Lusvarghi et al., 2021) blinded results of a study where 25 clinical stage SARS-CoV-2 antibody therapeutics (monoclonals, monoclonal cocktails and polyclonal/hyperimmunes) provided by various manufacturers were tested against a broad range of variants using a full length S protein HIV-based lentivirus pseudovirus assay (human 293T cell line stably expressing ACE2 and TMPRSS2 (293T-ACE2.TMPRSS2_s_)) to determine the effect of the mutations on the ability of the antibody therapeutics to neutralize the variants. The published results included neutralization activity against 60 pseudoviruses bearing spike protein mutations with single and multiple substitutions/deletions in several spike domains, including full set substitutions of the Alpha (B.1.1.7), Beta (B.1.351), Gamma (P.1), Epsilon (B.1.429), Iota (B.1.526), A.23.1, and R. 1 variants. Since this publication, results have been provided to the manufacturers on additional full sequence pseudovirus VOI/VOCs, including Delta/Delta+ (B.1.617.2), Kappa (B.1.617.1 V1+V2), and Lambda (C.37), among others. In addition, live virus results relative to the Washington isolate of SARS-CoV-2 (USA-WA1-2020, NR52281) for two Alpha (B.1.1.7) variant isolates, Beta (B.1.351), and the D614G variant results have been received from NIAID Integrated Research Facility and are provided in table 1 below for Emergent’s COVID-HIG.

## Discussion

Since the start of the COVID-19 outbreak several variants of SARS-CoV-2 have been identified with multiple amino acid substitutions in the spike protein that can enhance transmission, reduce/eliminate the effectiveness of neutralizing antibodies from natural infection, vaccination, or antibody therapeutics like convalescent plasma, monoclonals, and monoclonal cocktails (Andreano et al., 2020; Greaney et al., 2021; Tada et al., 2021; Wang et al., 2021c; Weisblum et al., 2020; West et al., 2021; Wibmer et al., 2021). These SARS-CoV-2 variants represent a significant long-term risk for high-risk patients especially those that don’t respond well to vaccination due to compromised immune systems. Antibody therapeutics that are resistant to losing potency to current and future variants are needed to help address these risks.

Emergent’s COVID-HIG (NP-028) polyclonal hyperimmune is currently being evaluated in a Ph 3 clinical study and has been shown to neutralize all 20 full sequence pseudovirus and 6 live virus variants tested in-vitro to date (table 1). COVID-HIG is manufactured using plasma from many individuals screened for high titers to SARS-CoV-2 and contains the combined broad range of polyclonal anti-SARS-CoV-2 antibodies from each plasma donor. This combined polyclonality has been shown to retain in-vitro neutralizing potency to SARS-CoV-2 variants compared to monoclonals/monoclonal cocktails as well as convalescent plasma.

Recently it has been shown (Schmidt et al., 2021) that individuals who were infected with SARS-CoV-2 and then vaccinated with mRNA vaccines produced significantly higher neutralizing antibody responses (compared to convalescent and vaccinated only). Plasma from convalescent individuals who were subsequently vaccinated with mRNA vaccines was also shown to be much more resistant to SARS-CoV-2 variants as well as a SARS-CoV-2 polymutant pseudovirus construct. The polymutant pseudovirus contained 20 spike protein substitutions (including 8 NTD and 8 RBD) selected from VOC as well as mutations that resulted from selective pressure by culturing the virus over several passages in the presence of convalescent plasma (details described in (Schmidt et al., 2021)). In that same publication, plasma from convalescent then mRNA vaccinated individuals was able to neutralize five SARS-CoV-2 variants (Alpha, Beta, Gamma, Delta, Iota) as well as SARS-CoV-1, two pangolin and two bat sarbecovirus pseudoviruses, whereas plasma from convalescent or vaccinated only donors had significantly lower or complete loss of neutralizing titers. The significantly higher neutralizing antibody titers and robust protection against current and potentially future SARS-CoV-2 variants seen with convalescent then mRNA vaccinated donor plasma provides a future opportunity for increased potency and even more robust variant resistance for Emergent’s COVID-HIG NP-028.

In conclusion, Emergent’s COVID-HIG (NP-028) has been shown to neutralize all virus variants tested to date and can adapt over time as new variants arise. The ability of Emergent’s COVID-HIG hyperimmune to maintain neutralizing activity across all VOC/VOIs tested to date has potential public health benefit due to the reduced neutralizing activity displayed by some monoclonal antibodies which could lead to diminished therapeutic efficacy as well as drive the creation of new escape mutants. In-vitro pseudovirus and live virus assays to SARS-CoV-2 variants are valuable tools to help evaluate the effectiveness of vaccines and antibody therapeutics to SARS-CoV-2 variants.

## Materials and Methods

### COVID-HIG (NP-028)

COVID-HIG is an investigational product that is not yet approved by the FDA and its safety and efficacy have not yet been established. COVID-HIG is a human hyperimmune product of purified immunoglobulin (IgG) fraction of human plasma containing antibodies to SARS-CoV-2. COVID-HIG is being evaluated for IM, SC or IV route of administration. COVID-HIG is prepared from pooled plasma collected at US Food and Drug Administration (FDA)-licensed and/or registered plasma and/or blood collection centers from healthy, adult donors who have elevated levels of SARS-CoV-2 antibodies. Plasma is collected from donors who meet all applicable requirements (e.g., FDA and Health Canada), for the type of plasma being collected. Plasma donors are carefully screened for eligibility by physical exam, and through questionnaires and interviews to assess risk of exposure to certain viruses or adventitious agents that may cause Creutzfeldt-Jakob disease (CJD) and its variant form (vCJD). Each plasma donation is tested for human immunodeficiency viruses 1/2 (HIV-1/2) and hepatitis C virus (HCV) antibodies, and as well as hepatitis B virus (HBV) surface antigen. Polymerase chain reaction tests for enveloped (HCV, HIV-1 and HBV) and non-enveloped [hepatitis A virus (HAV) and Parvovirus B-19] viruses are also performed on plasma minipools representing each of the individual donations used in COVID-HIG manufacturing. In addition, the manufacturing process for COVID-HIGIV contains orthogonal steps implemented specifically for virus clearance. The solvent and detergent step (using TnBP and TX-100, respectively) is effective in the inactivation of enveloped viruses such as human immunodeficiency virus (HIV), hepatitis B virus (HBV) and hepatitis C virus (HCV), among others. Virus filtration, using a Planova™20N virus filter, is effective for the removal of viruses based on their size, including both enveloped and non-enveloped viruses. These two viral clearance steps are designed to increase product safety by reducing the risk of transmission of enveloped and non-enveloped viruses. In addition to these two specific steps, the process step of anion-exchange chromatography was identified as contributing to the overall viral clearance capacity for small non-enveloped viruses. The COVID-HIG finished drug product is formulated with 250 mM proline and polysorbate 80 (PS80, 0.03% w/w) at target pH 5.8 and contains 10% protein (target 100 mg/mL; 10 g%) consisting of ≥90% IgG. COVID-HIG also may contain residual amounts of solvent (tri-n-butyl phosphate; TnBP) and detergent (Triton X-100®; TX-100) used to inactivate lipid-enveloped viruses during the manufacturing process. COVID-HIG activity against SARS-CoV-2 is determined by validated in-house potency assays (trimeric SARS-CoV-2 Spike protein antigen and Lentivirus neutralization) at product release and has been characterized for neutralizing potency by a wild-type SARS-CoV-2 assay (Bennett et al., 2021) as well

#### Production of SARS-CoV-2 Pseudovirus Variants and Neutralization Assay

Lentiviral pseudoviruses bearing Spikes of variants were generated and used for neutralization studies performed at FDB/CBER in stable 293T-ACE2/TMPRSS2 cells (BEI # NR-55351) as previously described (Neerukonda et al., 2021). Pseudoviruses with titers of approximately 106 relative luminescence units (RLU)/ml were incubated with four-fold serially diluted antibodies for two hours at 37°C prior to inoculation onto target cells in 96-well plates and scoring for luminescence activity 48 hours later. Titers were calculated using a nonlinear regression curve fit (GraphPad Prism software Inc., La Jolla, CA). The nAb concentration or inverse of the dilutions causing a 50% reduction of RLU compared to control (IC50) was reported as the neutralizing antibody titer. Antibodies not reaching 80% neutralization were considered resistant (R), and IC50 values were reported as greater than the highest concentration tested. The mean titer from at least two independent experiments each with intra-assay duplicates was reported as the final titer. WT was run as a control for each assay.

#### SARS-CoV-2 Live Virus Neutralization Assay

Live virus testing was done at NIAID/IRF (under BSL-3). Details of the method can be found in (Bennett et al., 2021). In brief, test material is diluted in a two-fold serial dilution through 12 steps in quadruplicate rows. A fixed amount of virus amounting to an MOI=0.5 is added to the diluted test material and incubated at 37°C for 1 hour. The test material + virus mixture is added to Vero E6 cells and incubated at 37°C/5% CO_2_ for 24 hours. Cells are fixed with formalin, removed from biocontainment, and probed with SARS-CoV/SARS-CoV-2 N protein specific mAb. Cells are probed with a fluorophore conjugated secondary antibody and counterstained with Hoechst nuclear dye. The number of SARS-CoV-2 positive cells is counted in four independent fields, each with >1000 cells, using a high content imaging system. The % positive wells relative to untreated controls is determined and the mean of four replicates per dilution is plotted against the concentration for each dilution. A four-parameter logistical analysis is performed to determine the 50% neutralization value. (Y=Bottom+(Top-Bottom)/(1+10^((LogEC50-X)*HillSlope)) (Prism, Graphpad). The SARS-CoV-2 Washington isolate is used as a control to determine fold reduction/increase in neutralization to each variant tested.

## Acknowledgements and disclosure of Potential Conflicts of Interest

Emergent is receiving funding from the U.S. Department of Defense’s Joint Program Executive Office for Chemical, Biological, Radiological and Nuclear Defense (JPEO-CBRND), in collaboration with the Defense Health Agency, under Other Transaction number W911QY-20-9-0013, and from the U.S. Department of Health and Human Services’ Biomedical Advanced Research Development Authority (BARDA) under Task Order HHSO100201200004I_75A50120F33006, for the development of COVID-HIG, a countermeasure to SARS-CoV-2. The U.S. Government is authorized to reproduce and distribute reprints for Governmental purposes notwithstanding any copyright notation thereon. The views and conclusions contained herein are those of the authors and should not be interpreted as necessarily representing the official policies or endorsements, either expressed or implied, of the U.S. Government. DT is an employee of Emergent BioSolutions Canada Inc.

## Limitations

Emergent’s COVID-HIG is currently in clinical trials and efficacy and safety against COVID-19 infection has yet to be established.

